# Association and impact of hypertension defined using the 2017 AHA/ACC guidelines on the risk of atrial fibrillation in the Atherosclerosis Risk in Communities Study

**DOI:** 10.1101/620393

**Authors:** Anas Rattani, J’Neka S. Claxton, Mohammed K. Ali, Lin Y. Chen, Elsayed Z. Soliman, Alonso Alvaro

## Abstract

Hypertension is an established risk factor for the development of atrial fibrillation (AF). We evaluated the association and population impact of hypertension, defined using the new 2017 guidelines, on risk of AF. In this analysis, we included 9,207 participants in the Atherosclerosis Risk in Communities study without history of cardiovascular disease or diabetes. Participants underwent blood pressure measurements at baseline and their antihypertensive medication use was assessed. AF was ascertained from study electrocardiograms, hospital records and death certificates. Cox proportional models were used to estimate hazard ratios (HR) and 95% confidence intervals (CI) of AF among individuals with hypertension based on the JNC7 and 2017 ACC/AHA guidelines. Poisson models were used to obtain risk ratios and calculate population-attributable fractions (PAFs). We identified 1,573 cases of incident AF during 22.1 years of mean follow-up. Prevalence of hypertension was 29% and 43% under the JNC7 and 2017 ACC/AHA definitions, respectively. HRs (95%CI) of AF in hypertensives versus non-hypertensives were 1.54 (1.39, 1.72) and 1.45 (1.31, 1.61) after multivariable adjustment under the old and new guidelines, respectively. The corresponding PAF (95%CI) using the old and new guidelines were 12% (9%, 14%) and 14% (10%, 18%), respectively. Overall, our analysis shows that even though the prevalence of hypertension using the new criteria is 50% higher than with the old criteria, this does not translate into meaningful increases in AF attributable to hypertension. These results suggest that prevention or treatment of hypertension based on the new (versus old) guidelines may have limited impact on AF incidence.

## Introduction

Atrial fibrillation (AF) is a common chronic arrhythmia, affecting between 2.7-6.1 million people in the United States.^1^ Common risk factors for AF include obesity, diabetes, smoking, heavy drinking, and hypertension.^2^ Among the risk factors listed, hypertension has the largest population attributable fraction for AF incidence and plays a major role in the management and prognosis of AF.^3–5^ Individuals with hypertension have a 1.7-fold higher risk of developing AF, with one in six cases of AF possibly due to hypertension.^3^ Hypertension is very common among individuals with AF, with studies showing prevalence of 69 to 90% of hypertension among AF patients.^4, 6, 7^ Thus, early detection and management of hypertension is key to preventing and managing AF.

The 7^th^ Joint National Committee (JNC7) defined hypertension as a systolic blood pressure (SBP) ≥140 mmHg or diastolic blood pressure (DBP) ≥90 mmHg, regardless of age. Blood pressure was divided into the following categories: Normal: SBP<120 and DBP<80, Prehypertension: SBP 120-139 or DBP 80-89, Stage 1 hypertension: SBP 140-159 or DBP 90-99, and Stage 2 hypertension: SBP≥160 or DBP≥100.^8, 9^ At the end of 2017, the American Heart Association/American College of Cardiology (AHA/ACC) released new guidelines lowering the threshold to define elevated blood pressure. The new recommended blood pressure diagnostic categories are: Normal: SBP<120 and DBP<80, Elevated: SBP 120-129 and DBP <80, Stage 1 hypertension: SBP 130-139 or DBP 80-89, Stage 2 hypertension: SBP ≥140 or DBP ≥90.^10, 11^ This change means more individuals will be diagnosed with hypertension. For example, an analysis of NHANES data comparing the 2014 and 2017 hypertension guidelines reported an increase in the prevalence of hypertension from 32 to 45%.^12^ However, it is uncertain whether individuals labeled as being hypertensive with the new guidelines are at similarly increased risk of AF, or whether the population impact of newly defined hypertension will have a similar impact in the incidence of AF.

The goal of this study was to evaluate the association between hypertension and risk of AF and the population attributable fraction of hypertension in AF using the new diagnostic categories in a prospective cohort free of cardiovascular disease and diabetes at baseline. Results from these analyses will contribute to inform the ideal blood pressure range for the prevention of AF as well as the potential population impact of preventing and treating hypertension under the new guidelines.

## Methods

### Study Population

For the present analysis, we used data from the Atherosclerosis Risk in Communities (ARIC) study cohort. The ARIC cohort aims to investigate the epidemiology of atherosclerosis, clinical atherosclerotic diseases, and variation in cardiovascular risk factors, treatment, and disease. The cohort study began in 1987, recruiting participants from four U.S. communities: Washington County in Maryland, Forsyth County in North Carolina, city of Jackson in Mississippi, and the northwest suburbs of Minneapolis in Minnesota. There were approximately 4,000 participants recruited from each community through probability sampling. A total of 15,792 participants aged 45-64 (7,082 were men and 11,526 were white) were enrolled. Detailed clinical, social, and demographic data were obtained at baseline in 1987-89. Participants have had additional evaluations in 1990-92, 1993-95, 1996-98, 2011-2013, and 2016-2017. The participants were also followed-up annually (biannually since 2012) by telephone to stay in contact, ascertain cardiovascular events, and to measure the health status of the cohort. More information about the design and objectives of the study can be found on the ARIC website as well as in published articles.^13^

For the present analysis, we included ARIC participants who had baseline blood pressure readings at visit 1 (1987-89). We excluded participants who had AF at baseline or missing ECG (N=346), individuals of a race other than white or black, as well as blacks from the Minneapolis and Washington County Centers (N=103), an eGFR value of less than 60 ml/min/1.73 m^2^ (N=73), and participants who had prevalent diabetes, coronary heart disease, stroke or heart failure (N=6,028). We also excluded participants who had missing values for the outcome, exposure, or covariates (N=35). After excluding participants who did not meet our study criteria, our final sample size was 9,207 participants (Figure 1 presents a flow chart for the final sample size).

**Figure 1:**
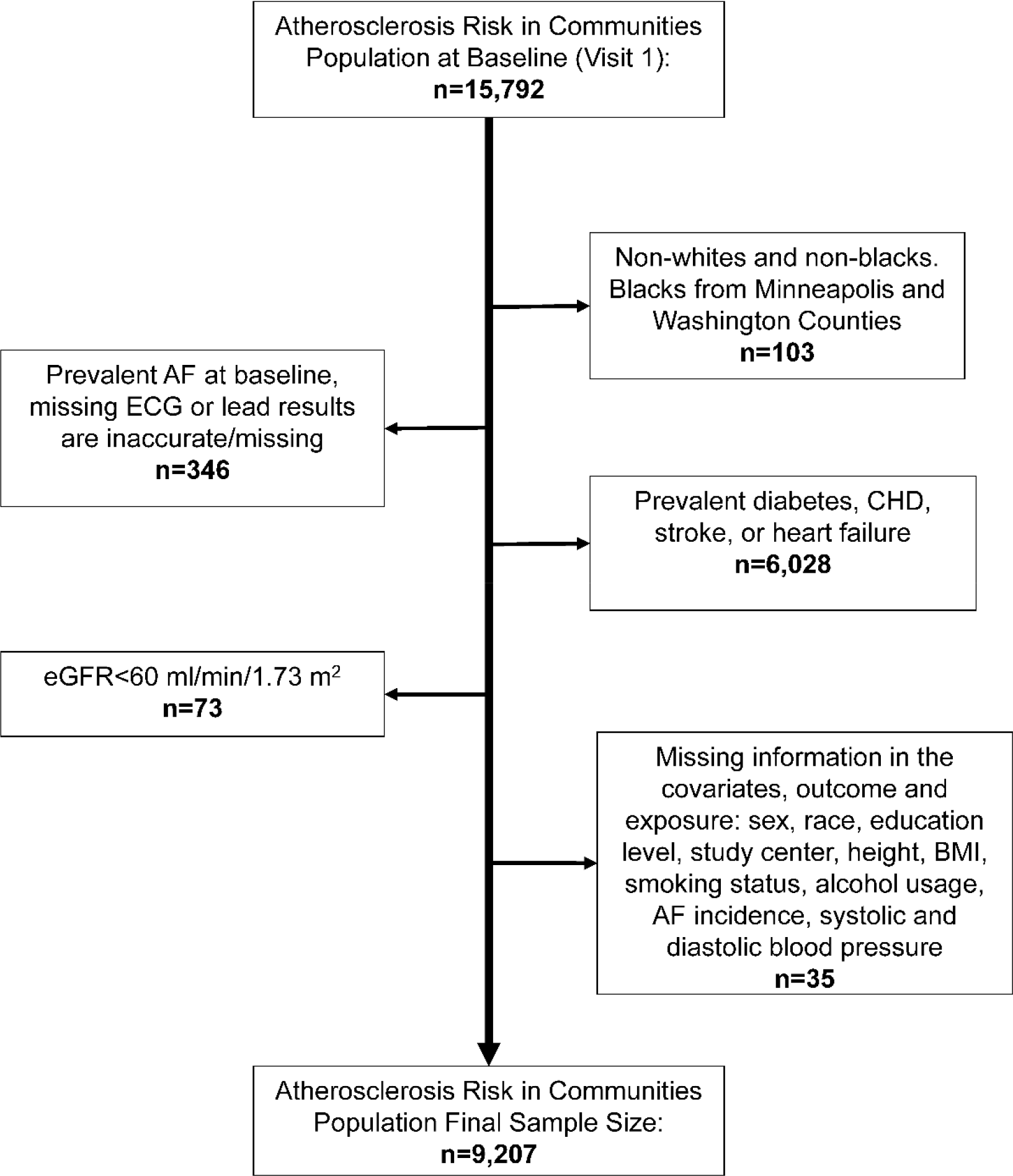
Flow Chart of Patients Excluded at Baseline, Atherosclerosis Risk in Communities Study, 1987.

### Assessment of blood pressure

At the baseline visit, sitting systolic and diastolic blood pressure was measured with a random zero sphygmomanometer 3 times at baseline after a 5-minute rest. The second and third measurements were averaged and used in the analysis. Use of blood pressure lowering medications was ascertained by asking participants to bring to the visit all medications they had been using over the previous 2 weeks. Baseline blood pressure and use of antihypertensive medication was used for all analyses.

### Assessment of Incident AF

We used three methods to identify cases of AF in the ARIC cohort: ECG performed at study visits, hospital discharge codes, and death certificates. The ECG studies were performed with a 12-lead ECG during study exams. The data obtained was transmitted electronically to the ARIC Central ECG Reading Center and processed using the GE Marquette 12-SL program. Presence of AF in the ECG was identified by a computer algorithm and then confirmed by a cardiologist. A cardiologist also read over ECGs with any other rhythm abnormalities to reduce the possibility of any missed AF incidents. Hospitalizations during the study period were identified with follow-up phone calls and monitoring local hospitals. Information such as discharge codes were collected from these hospitals by abstractors. If a participant had discharge codes ICD-9-CM codes 427.31 or 427.32 (ICD-10-CM code I48.x after October 1, 2015), then they were considered to have AF. Cases where a participant had open heart surgery in association with AF were excluded. Finally, if a patient had codes such as ICD-9 427.3 or ICD-10 I48 in their death certificates, then the participant was considered to have AF.^14^

### Assessment of covariates

Sex, race, education (categorized as grade school, high school but no degree, high school graduate, vocational school, college or graduate/professional school), smoking status (categorized as never, former or current smoker), and alcohol usage (categorized as never, former, current drinker) were obtained via self-report, while height and weight were measured with participants wearing light clothing. We calculated body mass index (BMI) as weight in kilograms divided by height in meters squared.

### Statistical Analysis

Analysis was conducted using SAS 9.4 statistical software. Cox proportional models were used to estimate the hazard ratios (HR) and 95% confidence interval (CI) of AF incidence among individuals with hypertension based on the JNC7 and 2017 AHA/ACC guidelines. For our independent variable, both the new and old hypertensive guidelines were divided into categories established by JNC7 and 2017 AHA/ACC, as indicated in Table 1. Participants using antihypertensive medication were labeled as having hypertension (JNC7) or stage 2 hypertension (2017 AHA/ACC) regardless of their visit blood pressure.

**Table 1:** Categories of Blood Pressure Established by JNC7 and 2017 ACC/AHA Guidelines.

| **JNC 7** |  |  |  | **2017 ACC/AHA** |  |  |  |
| --- | --- | --- | --- | --- | --- | --- | --- |
|  | **SBP** |  | **DBP** |  | **SBP** |  | **DBP** |
| Normal | <120 | and | <80 | Normal | <120 | and | <80 |
| Prehypertension | 120-139 | or | 80-90 | Elevated | 120-129 | and | <80 |
| Hypertension | ≥140 | or | ≥90 | Stage 1 | 130-139 | or | 80-89 |
|  |  |  |  | Stage 2 | ≥140 | or | ≥90 |

SBP/DBP measurements are in mmHg.
JNC 7: 7th Joint National Committee, ACC/ AHA: American College of Cardiology/American Heart Association, SBP: systolic blood pressure, DBP: diastolic blood pressure

Two separate analyses were conducted to fully characterize the impact of changing the definition of hypertension. The first analysis considered hypertension as a binary variable. For the JNC7 guideline, prehypertension and normal were combined in the reference group. For the 2017 AHA/ACC guideline, elevated and normal blood pressure were combined as the reference group whereas stage 1 and stage 2 were combined to define hypertension. We conducted a stratified analysis by sex and race to explore effect modification. The second analysis considered hypertension classified into more specific categories in both the JNC7 and 2017 AHA/ACC guidelines. Using a normal blood pressure as the reference in both guidelines, we calculated HRs of AF among individuals with prehypertension/elevated, hypertension, stage 1, or stage 2. Covariate adjustment was done through two separate models. Model 1 adjusted for age, sex, and race, while model 2 additionally adjusted for education, study center, height, BMI, smoking status, and alcohol use.

We calculated population-attributable fractions (PAFs) of AF by hypertension categories to determine the possible impact of preventing hypertension on AF occurrence. PAFs were computed according to the following formula: PAF=pd_i_[(RR_i_-1)/RR_i_], where pd_i_ is the proportion of cases falling into *i*th exposure level and RR_i_ is the relative risk (RR) comparing *i*th exposure level with unexposed group (i=0).^15^ Poisson models were used to estimate RRs. The offset in the Poisson model was calculated as the natural logarithm of the time from visit 1 to AF incidence, death or lost to follow up until December 31, 2015, whichever came first. Ninety five percent confidence intervals (95%CI) for the PAF were obtained applying the corresponding 95%CI of the RR to the PAF formula above.

## Results

### Basic Demographic Characteristics of participants in ARIC study

We included 9,207 eligible adults in our final sample. The mean age at baseline was 53.7 years old (SD=5.1). The sample was 76% white and 56% women. The percentage of individuals taking hypertension medication in the cohort was 19%. Based on the JNC7 guidelines, 46% individuals had normal blood pressure, 25% of individuals were prehypertensive and 29% of individuals were hypertensive (Table 2). Based on the 2017 ACC/AHA guidelines, 46% individuals had normal blood pressure, 11% of individuals had elevated blood pressure, 14% of individuals were stage 1 hypertensive, and 29% of individuals were stage 2 hypertensive (Table 2).

**Table 2:** Demographic Characteristics of Participants in the ARIC Study Based on the JNC 7 and ACC/AHA Guideline Categories.

| JNC7 Categories | Normal | Prehypertension |  | Hypertension |
| --- | --- | --- | --- | --- |
| N (%) | 4273 (46.4) | 2315 (25.1) |  | 2619 (28.5) |
| Age (years) | 52.5 (5.5) | 54.3 (5.7) |  | 55.0 (5.6) |
| White | 3709 (86.8) | 1731 (74.8) |  | 1559 (59.5) |
| Women | 2532 (59.3) | 1180 (51.0) |  | 1471 (56.2) |
| Completed high school | 3665 (85.8) | 1844 (79.7) |  | 1877 (71.7) |
| Ever smokers | 2519 (59) | 1325 (57.3) |  | 1481 (56.5) |
| Ever drinkers | 3433 (80.3) | 1733 (74.9) |  | 1876 (71.8) |
| BMI (kg/m2) | 25.8 (4.2) | 27.6 (5.2) |  | 28.7 (5.8) |
| Height (cm) | 168.5 (9.3) | 169.1 (9.5) |  | 168.2 (9.3) |
| SBP | 106.2 (8.3) | 126.1 (6.4) |  | 136.6 (20.1) |
| DBP | 66.4 (7.1) | 76.4 (7.5) |  | 81.6 (11.8) |
| Hypertension medication | -- | -- |  | 1732 (66.2) |

| ACC/AHA categories | Normal | Elevated | Stage 1 hypertension | Stage 2 hypertension |
| --- | --- | --- | --- | --- |
| N (%) | 4273 (46.4) | 986 (10.7) | 1329 (14.4) | 2619 (28.5) |
| Age (years) | 52.5 (5.5) | 55.2 (5.7) | 53.6 (5.7) | 55.0 (5.6) |
| White | 3709 (86.8) | 797 (80.8) | 934 (70.3) | 1559 (59.5) |
| Women | 2532 (59.3) | 529 (53.7) | 651 (49.0) | 1471 (56.2) |
| Completed high school | 3665 (85.8) | 784 (79.5) | 1060 (79.8) | 1877 (71.7) |
| Ever smokers | 2519 (59) | 580 (58.9) | 745 (56) | 1481 (56.5) |
| Ever drinkers | 3433 (80.3) | 745 (75.5) | 988 (74.4) | 1876 (71.8) |
| BMI (kg/m2) | 25.8 (4.2) | 27.2 (4.8) | 27.9 (5.3) | 28.7 (5.8) |
| Height (cm) | 168.5 (9.3) | 168.5 (9.5) | 169.6 (9.4) | 168.2 (9.3) |
| SBP | 106.2 (8.3) | 123.9 (2.8) | 127.8 (7.8) | 136.6 (20.1) |
| DBP | 66.4 (7.1) | 71.5 (6.2) | 80.0 (6.2) | 81.6 (11.8) |
| Hypertension medication | -- | -- | -- | 1732 (66.2) |

Numbers correspond to mean (SD) and N (percentages).
JNC 7: 7th Joint National Committee, ACC/ AHA: American College of Cardiology/American Heart Association, SBP: systolic blood pressure, DBP: diastolic blood pressure

### Association of hypertension (binary) with AF incidence using JNC 7 and 2017 ACC/AHA definitions

During a mean follow-up of 22.1 years, we identified 1,573 cases of incident AF overall. Using the JNC7 definition, the incidence rate of AF per 1000 person-years were 6.6 and 10.8 for no hypertension and hypertension respectively. The HR of AF in hypertension compared to no hypertension was 1.65 (95% CI 1.49, 1.83) after adjusting for age, sex and race, and 1.54 (95%CI 1.39, 1.72) after multivariable adjustment. Corresponding AF rates using the 2017 AHA/ACC guidelines definition were 6.4 and 9.6 per 1000 person-years for no hypertension and hypertension, respectively. The HR of AF was 1.52, (95% CI 1.37, 1.68) after adjusting for age, sex and race and 1.45, (95%CI 1.31, 1.61) after multivariable adjustment (Table 3).

**Table 3:** Hazard Radios (95% Confidence intervals) of Atrial Fibrillation According to Hypertension Defined According to JNC 7 and 2017 ACC/AHA Guidelines, ARIC 1987-2015.

| **JNC 7** | **No hypertension** | **Hypertension** |
| --- | --- | --- |
| **No. of AF cases** | 992 | 581 |
| **Person-years** | 149742 | 53910 |
| **Incidence rate (per 1000 PY)** | 6.6 | 10.8 |
| **Model 1 [HR (95%CI)]** | 1 (ref.) | 1.65 (1.49, 1.83) |
| **Model 2 [HR (95%CI)]** | 1 (ref.) | 1.54 (1.39, 1.72) |
| **2017 ACC/AHA** | **No hypertension** | **Hypertension** |
| **No. of AF cases** | 770 | 803 |
| **Person-years** | 120370 | 83283 |
| **Incidence rate (per 1000 PY)** | 6.4 | 9.6 |
| **Model 1 [HR (95%CI)]** | 1 (ref.) | 1.52 (1.37, 1.68) |
| **Model 2 [HR (95%CI)]** | 1 (ref.) | 1.45 (1.31, 1.61) |

Model 1: Adjusted for age, sex, and race.
Model 2: Adjusted for age, sex, race, height, education, field center, body mass index, smoking and drinking status
JNC 7: 7th Joint National Committee, ACC/ AHA: American College of Cardiology/American Heart Association, PY: person-years

In analyses stratified by sex and race, association of hypertension, as defined by both JNC7 and 2017 AHA/ACC guidelines, were similar across groups, with the exception of a significant interaction by sex in the 2017 AHA/ACC guidelines (p=0.02) (Supplementary Table I). Hypertension was more strongly associated with AF incidence in women (HR 1.68, 95%CI 1.44, 1.96) than in men (HR 1.27, 95%CI 1.10, 1.47).

### Association of blood pressure with AF using JNC 7 and 2017 ACC/AHA guideline categories

Using the JNC7 categories, the AF incidence rates per 1000 person-years were 5.9, 8.0, and 10.8 for normal blood pressure, prehypertension, and hypertension, respectively. The HRs (95%CI) of AF for prehypertension and hypertension, compared to normal blood pressure, were 1.23 (1.08, 1.40) and 1.69 (1.50, 1.92) respectively after multivariable adjustment. Using 2017 AHA/ACC guideline categories, the incidence rates for AF per 1000 person-years were 5.9, 9.2, 7.6, and 10.8 for normal, elevated, stage 1, and stage 2, respectively. The HR (95%CI) for elevated, stage 1 and stage 2, compared to normal blood pressure, were: 1.22 (1.04, 1.45), 1.24 (1.06, 1.45), and 1.69 (1.50, 1.92), respectively, after multivariable adjustment (Table 4).

**Table 4:** Hazard Ratios (95% Confidence Intervals) of Atrial Fibrillation by Categories of Blood Pressure According to JNC 7 and 2017 ACC/AHA Definitions, ARIC 1987-2015.

| JNC 7 | Normal | Prehypertension | Hypertension |  |
| --- | --- | --- | --- | --- |
| No. of AF cases | 584 | 408 | 581 |  |
| Person-years | 98994 | 50748 | 53910 |  |
| Incidence rate (per 1000 PY) | 5.9 | 8.0 | 10.8 |  |
| Model 1 [HR (95%CI)] | 1 (ref.) | 1.26 (1.11, 1.43) | 1.82 (1.61, 2.05) |  |
| Model 2 [HR (95%CI)] | 1 (ref.) | 1.23 (1.08, 1.40) | 1.69 (1.50, 1.92) |  |
| 2017 ACC/AHA | Normal | Elevated | Stage 1 | Stage 2 |
| No. of AF cases | 584 | 186 | 222 | 581 |
| Person-years | 98994 | 20280 | 29372 | 53910 |
| Incidence rate (per 1000 PY) | 5.9 | 9.2 | 7.6 | 10.8 |
| Model 1 [HR (95%CI)] | 1 (ref.) | 1.28 (1.08, 1.51) | 1.25 (1.07, 1.46) | 1.82 (1.61, 2.05) |
| Model 2 [HR (95%CI)] | 1 (ref.) | 1.22 (1.04, 1.45) | 1.24 (1.06, 1.45) | 1.69 (1.50, 1.92) |

Model 1: Adjusted for age, sex, and race.
Model 2: Adjusted for age, sex, race, height, education, field center, body mass index, smoking and drinking status
JNC 7: 7th Joint National Committee, ACC/ AHA: American College of Cardiology/American Heart Association, PY: person-years,

### Population Attributable Fraction of the JNC7 and 2017 AHA/ACC guidelines

Using the JNC 7 guidelines, the PAFs were 3% (95% CI [0, 6]) for prehypertension and 11% (95% CI [7, 14]) for hypertension. In contrast, using categories defined in the 2017 ACC/AHA guidelines, the PAFs were 2% (95% CI [0, 3]), 1% (95% CI [-1, 3]), and 11% (95% CI [7, 14]) for elevated, stage 1 and stage 2 hypertension, respectively (Table 5). When the data was recategorized into binary variables, the prevalence of hypertension was 29% using the JNC 7 definition and 43% with the 2017 ACC/AHA definition. The PAF was 12% (95% CI [9, 14]) and 14% (95% CI [10, 18]) under the old and new guidelines respectively (Table 6).

**Table 5:** Rate Ratios and Population Attributable Factor of Atrial Fibrillation by Blood Pressure Categories According to JNC 7 and 2017 ACC/AHA Guidelines, ARIC 1987-2015.

| JNC 7 | Normal |  | Prehypertension |  | Hypertension |
| --- | --- | --- | --- | --- | --- |
| Prevalence, % | 46.4 |  | 25.1 |  | 28.5 |
| RR (95 %CI)* | 1 (ref.) |  | 1.13 (0.99,1.28) |  | 1.40 (1.24, 1.59) |
| PAF % (95% CI) |  |  | 3 (0, 6) |  | 11 (7, 14) |
| 2017 ACC/AHA | Normal |  | Elevated | Stage 1 | Stage 2 |
| Prevalence, % | 46.4 |  | 10.7 | 14.4 | 28.5 |
| RR (95% CI)* | 1 (ref.) |  | 1.16 (0.98, 1.37) | 1.10 (0.94,1.29) | 1.40 (1.06, 1.24) |
| PAF % (95% CI) |  |  | 2 (0, 3) | 1 (-1, 3) | 11 (7, 14) |

\* Adjusted for age, sex, race, height, education, field center, body mass index, smoking, drinking
JNC 7: 7th Joint National Committee, ACC/ AHA: American College of Cardiology/American Heart Association, RR: Rate Ratios, and PAF: Population Attributable Factor

**Table 6:**
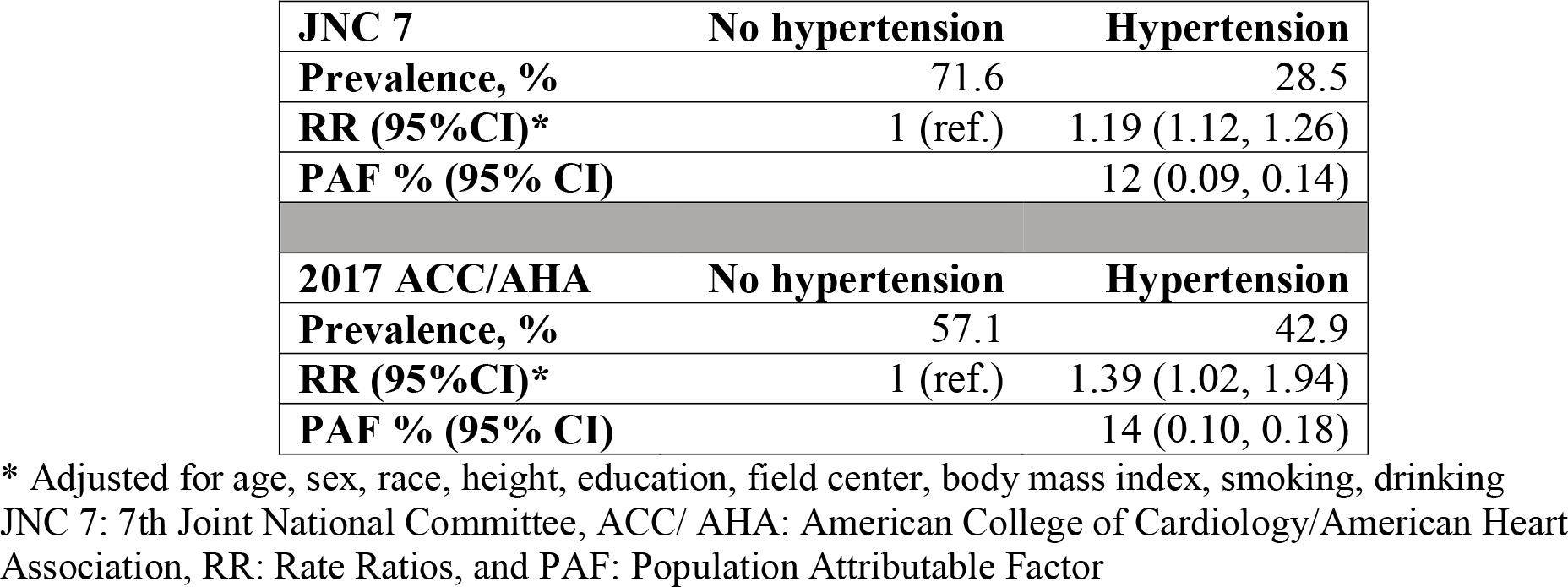
Rate Ratios and Population Attributable Factor of Atrial Fibrillation by Hypertension Definition According to JNC 7 and 2017 ACC/AHA Guidelines, ARIC 1987-2015.

## Discussion

In this analysis of a large community-based cohort without cardiovascular disease or diabetes at baseline, we found that the relative risk of AF associated with hypertension was similar using the JNC7 and 2017 ACC/AHA definitions. Also, despite an almost 50% increase in the prevalence of hypertension using the new 2017 ACC/AHA definition compared to JNC7 (from 29% to 43%), there was no excess risk of AF and the PAF of AF only increased from 12% to 14%. These results suggest that the extended definition of hypertension based in the guidelines may have a limited additional impact in the prevention of AF.

Hypertension is very common among individuals with AF, both conditions frequently coexisting. It has the largest population attributable fraction for AF incidence and is an important focus in the management and prognosis of AF.^3–5^ Studies have also shown that once hypertension occurs, an individual is predisposed to developing AF even if the blood pressure improves in later years.^3^ Thus, understanding the risk of AF in association with hypertension is crucial in preventing AF, reducing AF incidence rates, and subsequently preventing strokes. Several guidelines have been released over the years to identify individuals with hypertension based on an increased risk of adverse outcomes and prevent its deleterious consequences by keeping blood pressure at optimal levels. The most recent hypertension guideline, released in the fall of 2017 by the ACC/AHA, refined the guidelines released by JNC7 and JNC8 by lowering the threshold to define hypertension. Consequently, more individuals are diagnosed with hypertension under the new guidelines. The rationale for this change is based on the observed increased risk of cardiovascular disease among individuals in the JNC7 prehypertensive category and results from the SPRINT trial, showing cardiovascular benefit in the treatment of blood pressure, targeting a SBP of <120 mmHg.^16^ However, the risk of AF among individuals diagnosed with hypertension using the new guidelines is uncertain. Our results suggest that the risk of AF in participants with stage 1 hypertension according to the 2017 guidelines (part of the prehypertension category using the JNC7 definition) is only slightly increased compared to normotensive individuals.

We estimated that the PAF for AF from hypertension using the new guidelines barely increased compared to hypertension defined using the JNC7 definition. A prior analysis of the ARIC population showed that borderline blood pressure levels (SBP 120 - 139 mmHg or DBP 80 – 89) explained an additional 3% of AF cases. In that prior ARIC analysis, PAF from hypertension was 22% of incident AF and this number increased to 24% adding borderline levels of blood pressure,^5^ consistent with our new results. Discrepancies in the overall PAF from hypertension could be explained by the exclusion of persons with prevalent cardiovascular disease in this analysis.

### Strengths and Limitations

Our study had important strengths. First, we had a large sample size and long follow-up. The study participants were from four geographically diverse communities and the final sample size included 9,207 individuals, with a total of 1,573 AF events, providing enough events in each category. Additionally, our study had extensive information on other risk factors for AF, allowing us to adjust for potential confounders. However, there were several limitations in our study. First, though we adjusted for major potential confounders, other common causes of hypertension and AF may have biased the results. Secondly, the study did not differentiate between AF subtypes such as paroxysmal, persistent, or permanent AF. Thirdly, some cases of AF may have been missed due to the method of AF ascertainment, which relies on hospital discharge codes, ECGs conducted at study visits, and death certificates. Thus, paroxysmal and asymptomatic AF cases would have been less likely to be identified. Finally, we only considered baseline blood pressure measurement and, therefore, our analysis does not evaluate the impact of trajectories of blood pressure over time, which are known to impact AF risk.^17^

In conclusion, our study showed that hypertension defined using the 2017 AAC/AHA guidelines only led to slight increases in PAF values. These results indicate that the new hypertension guidelines may have a limited impact in the burden of AF in the population. Additional studies are needed to confirm these observations.

## Supporting information

Supplementary Results

## Acknowledgements

The authors thank the staff and participants of the ARIC study for their important contributions.

## Source of funding

The Atherosclerosis Risk in Communities study has been funded in whole or in part with Federal funds from the National Heart, Lung, and Blood Institute, National Institutes of Health, Department of Health and Human Services, under Contract nos. (HHSN268201700001I, HHSN268201700002I, HHSN268201700003I, HHSN268201700005I, HHSN268201700004I).

This work was supported by American Heart Association grant 16EIA26410001 (Alonso).

## Conflict of interest

None

