## Supplementary Results for "Association and impact of hypertension defined using the 2017 AHA/ACC guidelines on the risk of atrial fibrillation in the Atherosclerosis Risk in Communities Study"

**Supplementary Table I: Hazard Ratios (95% Confidence Intervals) of Atrial Fibrillation by Hypertension Definitions Stratified by Race and Sex, ARIC 1987-2015**

| **JNC 7** | **Hypertension** |
| --- | --- |
| **Women** | 1.64 (1.41, 1.92) |
| **Men** | 1.46 (1.26, 1.70) |
| **p-value for interaction** | 0.41 |
| **Whites** | 1.49 (1.32, 1.69) |
| **Blacks** | 1.81 (1.40, 2.34) |
| **p-value for interaction** | 0.18 |
| **2017 ACC/AHA** | **Hypertension** |
| **Women** | 1.68 (1.44, 1.96) |
| **Men** | 1.27 (1.10, 1.47) |
| **p-value for interaction** | 0.02 |
| **Whites** | 1.45 (1.30, 1.63) |
| **Blacks** | 1.49 (1.11, 1.99) |
| **p-value for interaction** | 0.79 |

Adjusted Age, sex, race, height, education, field center, body mass index, smoking and drinking status. Hypertension according to JNC7 defined as systolic blood pressure ≥140 mmHg or diastolic blood pressure ≥90 mmHg or use of antihypertensive medication. Hypertension according to 2017 ACC/AHA defined as systolic blood pressure ≥130 mmHg or diastolic blood pressure ≥80 mmHg or use of antihypertensive medication.
